# Validation and characterization of a viral antisense transcript with Northern blot analysis and qRT-PCR

**DOI:** 10.1101/2023.01.18.524592

**Authors:** Mehmet Kara

## Abstract

The transcription of mammalian genomes has been shown to possess an intriguing complexity and numerous novel RNA molecules have been identified in the last 10-15 years. Viruses with large DNA genomes, especially herpesviruses, are also shown to generate many different RNA species and some of them may function as long non-coding RNAs. Viral genomes harbor several genes within close proximity to each other and can generate multigenic transcripts in addition to commonly observed antisense transcription. It is essential to study the biological roles of these transcripts aside from the protein-coding counterparts. A transcriptionally complex locus can be studied with a combination of methods. Generally, quantitative PCR assays are very commonly used for expression analyses of the transcripts of interest. Here an example from a gammaherpesvirus is discussed in more detail. A recently developed method, for the resolution of complicated transcriptomes for viral genomes, elucidated multiple antisense transcripts from ORF63-64 locus in murine gammaherpesvirus 68 (MHV68). In order to identify the roles of these new transcripts, quantitative PCR assays may not be enough alone and should be supported by alternative methods such as Northern blots. A more detailed transcriptional map of the locus of interest is useful to design experimental strategies and perform functional studies.

## 1. Introduction

Over the last decade, developments and the relatively cheap availability of new sequencing technologies have been greatly increased. These rapid advancements caused a paradigm shift in our understanding of how much of the mammalian genome is being actively transcribed (Derrien et al., 2012; Djebali et al., 2012; Khalil et al., 2009). Through enormous transcriptomic data analysis, a vast array of intergenic, intronic, antisense, alternatively spliced, and alternative promoter-using transcripts has been identified. As opposed to the protein coding transcripts, these long, mostly 5’ capped and polyadenylated transcripts are not associated with the cellular translational machinery therefore; named long non-coding RNAs (lncRNA) (Ulitsky & Bartel, 2013). Interestingly, majority of the non-coding RNAs reside in the nucleus, which is consistent with the idea of these RNA molecules are not translated and they may function through sequence specific interactions with DNA and/or secondary motifs with proteins (Guh et al., 2020). Therefore characterization of these recently identified transcripts is important for functional studies and understanding how they perform specific key roles in cellular processes.

Viruses, with their limited capacity genomes, usually contain overlapping and bidirectional genes (Schlub & Holmes, 2020). They have been evolved to use similar transcriptional strategies like their host to produce many different coding and non-coding RNA molecules (Boldogkői et al., 2019; Tombácz et al., 2017) and it poses a difficult challenge to characterize and study the function of these recently identified transcripts. Even though sequencing information from long RNA molecules by PacBio’s SMRT (Single Molecule Real Time) technology combined with short Illumina based sequencing data have provided us high resolution annotation for the transcriptional complexity of the dense viral genomes (O’Grady et al., 2016), it is still essential to validate the novel transcripts experimentally. A given locus in viruses, as indicated by this study and also mammalian genomes to some extent, have the ability to generate several transcripts which are not depicted by the standard repositories such as NCBI.

Herpesviruses have dsDNA genomes ranging from 100 to 250 kilobases in size and can code for up to 235 proteins (Arvin et al., 2007). Recently many viruses in this family has been shown to generate putative lncRNAs during lytic and/or latent life cycle (Arias et al., 2014; Canny et al., 2014; O’Grady et al., 2016, 2019). In order to study the function of these transcripts, one of mainly used assays in the laboratory is real time PCR (qPCR) assays (Elfman & Li, 2020; Shi et al., 2020; Su & Lai, 2018). In a standard qPCR test, an RNA sample is converted to cDNA mainly by the use of random priming and PCR amplified by using primers 100-200 nucleotides apart from each other. Therefore, it becomes impossible to differentiate between the distinct isomeric forms as well as the antisense transcripts in terms of quantification of the desired gene product. The second part of this problem can be addressed by utilization of gene specific primers for the initial cDNA conversion however, different isomeric forms can still be captured in the qPCR assay. Therefore, it is beneficial to have a transcriptional map of given locus by strand specific Northern blot and then, distinct transcripts can be studied in depth. While examining the overlapping regions in the mammalian genome, a similar problem may arise. Therefore, even though the short read sequencing data yields an incredible amount of information and the regions of interest are evaluated with qRT-PCR assays, the concept of the alternative transcript isoforms and antisense transcription should not be overlooked.

Murine gammaherpesvirus 68 (MHV68) provides a valuable model system to study human or related gammaherpesvirus pathology. The transcriptional schematic of the virus has been studied over the years (Canny et al., 2014; Cheng et al., 2012; Virgin et al., 1999). The most recent annotation of viral transcriptome is published and data is available (O’Grady et al., 2019). In that study, a bioinformatics pipeline named TRIMD (<u>T</u>ranscriptome <u>R</u>esolution through Integration of <u>M</u>ultiplatform <u>D</u>ata), is utilized to identify ∼250 novel transcripts during lytic MHV68 infection. TRIMD pipeline uses PacBio long read sequencing, Illumina sequencing and 5’ cap sequencing (CAGE) data for RNA isoform identification. The antisense transcripts in this paper is partially discussed as a part of Northern blot protocol publication for a method paper (Kara & Tibbetts, 2021). Here in this work, a further examination with different probes and primers to map the transcriptome is provided. Additionally, a time course for the transcripts and latency expression profiles are analyzed.

## 2. Materials and methods

### 2.1 Cell culture and virus infections

For the cell culture and infections the mouse fibroblast cell line NIH 3T12 was used. Cells were maintained in Dulbecco’s modified Eagle’s medium (DMEM) with 10% fetal calf serum with antibiotics at 100 U/mL of penicillin and streptomycin concentration and 2 mM L-glutamine. MHV68.OR73Bla, a recombinant virus which expresses beta lactamase as a fusion to the LANA protein encoded by ORF73 (Nealy et al., 2010), was used for the viral infections. A20 mouse B cell lymphoma cell line and its derivative A20-HE.2.1(Forrest & Speck, 2008) which carries viral genome under antibiotic selection is used for latency model. B cell lines were maintained in RPMI medium with the same conditions above in addition to 50 µM β-mercaptoethanol. A20-HE.2.1 cell line kept under 300 µg/ml hygromycin B for viral genome persistence.

### 2.2 RNA extraction and Northern blot analysis

RNA extraction and Northern blot protocol is mainly based on the previously published protocol (Kara & Tibbetts, 2021). Briefly, fibroblasts were infected with 5 plaque forming unit MHV68 virus per cell in a 10 cm cell culture plate. At designated time points post infection, cells were washed with PBS and collected into 1 mL Trizol. 0.2 mL of chloroform was added to the Trizol sample, the upper phase was collected, then precipitated with isopropanol and washed with ice-cold 75% ethanol. Total RNA was resuspended in DEPC-treated deionized water and kept at -80°C.

For the northern blot protocol, around 3 to 5 μg total RNA from samples were loaded onto a 6% formaldehyde-containing 1% agarose gel in the fume hood with RNA Millenium Marker (Ambion) as RNA ladder. The gel was run ∼80 V for 3-4 hours in MOPS (3-(N-morpholino)propanesulfonic acid) buffer, then blotted onto a Hybond XL nylon membrane (Life Technologies) overnight with Turbolotter kit in 20X SSC (Standard Saline Citrate) buffer. The membrane was washed, RNA was crosslinked to the membrane by UV, and membrane was stained with 0.02% methylene blue for visualization of ribosomal RNA integrity and markers. The crosslinked membrane was prehybridized at 62°C for 2-4 hours in ULTRAhyb (Ambion) buffer and then labeled probe was added for overnight incubation. The next day the membrane was washed once with the each of the following buffers; 2x SSC, 1x SSC and 0.1X SSC with 0.1% SDS and exposed to a film at 80°C for appropriate time.

The radioactively labeled riboprobes were prepared with Maxiscript T7/Sp6 (Thermofisher). Briefly, the probe template was PCR amplified with primers containing T7 and Sp6 promoters on each side (given in the Table). The template was labeled with the kit by adding 10 μCi alpha-CTP (Perkin Elmer) to the reaction for four hours at 37°C. The DNA template was digested with DNase for 20 minutes, then the reaction was stopped by adding EDTA. The probe was used without purification.

**Table.**
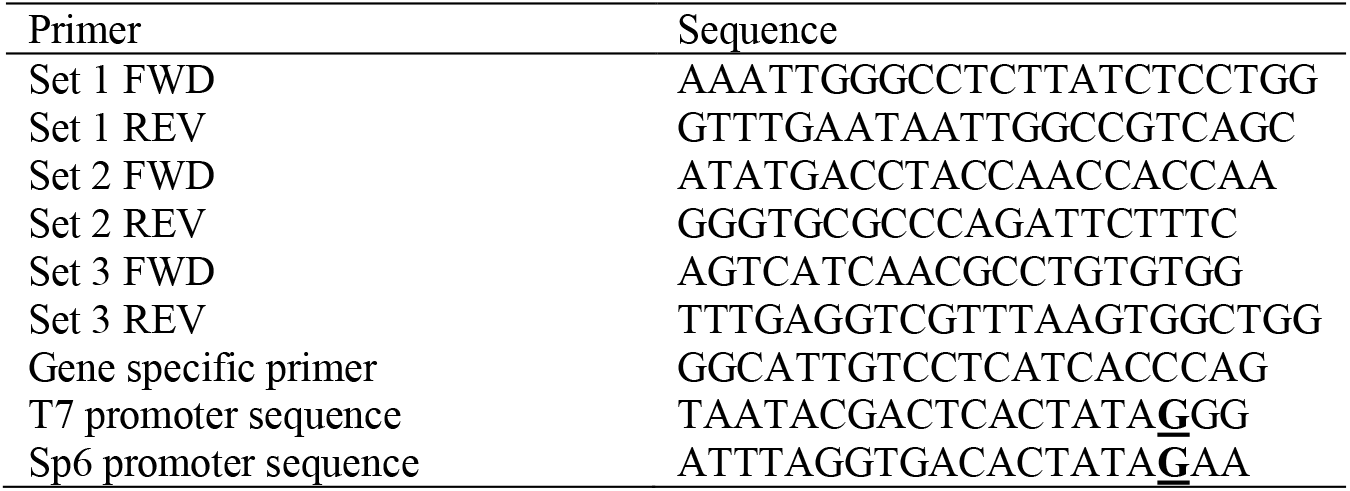
Primer list for qPCR and Northern blots. T7 or Sp6 promoter sequence containing primers are ordered with the same sequence for Set 1, 2, 3. The underlined bold G nucleotide in the promoter sequence is the first nucleotide of the RNA probe.

### 2.3 Quantitative reverse transcription (qRT) PCR

2μg of total RNA is converted to cDNA by an initial DNase digestion with Turbo DNase (Ambion) following either random priming or gene specific primers by iScript cDNA synthesis kit (BioRad). 20 μL of cDNA reaction is diluted 5 times before the quantitative PCR reaction. 2 μL of the cDNA sample is used for each triplicate of PCR reaction. 2X Master Mix (Fermentas) with the primers listed in the Table is used for qPCR amplification in CFX96 BioRad Real Time PCR machine. Data were analyzed with Graphpad Prism software.

### 2.4 Integrative Genomics Viewer data visualization

Original sequencing data is published and available at <u>GSE117651</u> from NCBI. The TRIMD analysis package can be obtained https://github.com/flemingtonlab/TRIMD and the Integrative Genomics Viewer (IGV) is downloadable from https://software.broadinstitute.org/software/igv/. The data sets can be downloaded and analyzed free of charge. Briefly, using the MHV68 genome as reference the .bed files are displayed in IGV and 14 kb region spanning ORF63-64 antisense transcript is visualized.

## 3. Results

### 3.1 Visualization of the viral transcripts in Integrative Genome Viewer (IGV)

A 14 kb region harboring the antisense transcripts, which stem from the ORF63-64 locus, is depicted in **Figure 1**. The region contains 17 total transcripts validated with TRIMD analysis. Majority of the transcripts have scores lower than 10. The score can be roughly interpreted as lower expression and detection with the method. Some of the transcripts have splice junctions and none of the spliced transcripts have scores higher than 7. One interesting observation is that, even though ORF64 is annotated in NCBI, the RNA product of this gene is not detected by this sequencing analysis. ORF64 mRNA should be a rightward transcript as opposed to the antisense transcripts synthesized from negative strand and going leftwards. The locations of the primers and probes used in this study is depicted in **Figure 1** and 5 major transcript (with scores higher than 50) can be expected from the locus.

**Figure 1.**
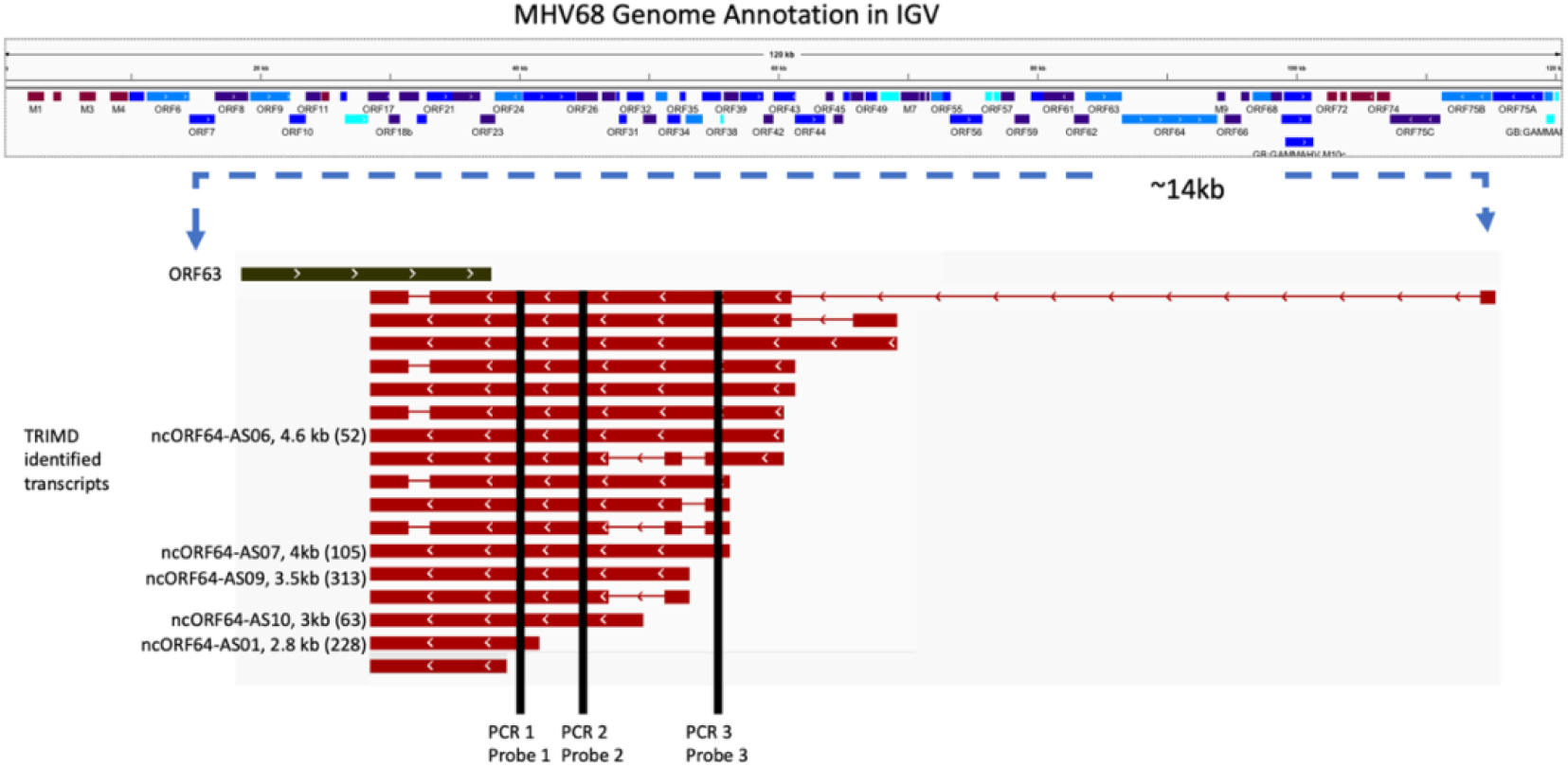
IGV visualization of ORF6364 antisense transcripts and locations of the qPCR regions and probes for Northern blots. The name and length is given next to the transcript line. Arrows indicate the transcriptional direction. Numbers in parentheses are TRIMD scores.

### 3.2 Northern blot analysis

RNA samples from mock or infected cells are used to detect the transcriptional complexity from the region. 3 sets of radiolabeled strand specific probes are generated and used as shown in **Figure 1**. According to the TRIMD analysis, the major transcripts should be around 1.9 kb, 3 kb, 3.5 kb, 4 kb and 4.6 kb in length. Probe 1 detected 5 major transcripts which are around 2 kb, 3 kb, 4 kb, 4.5 kb and an RNA molecule larger 9 kb and its size cannot be determined correctly by the RNA ladder used here (**Figure 2**). In general, the transcripts are in very close size range for the sequencing data identified transcripts. Interestingly, the large RNA was not detected in the TRIMD analysis. This is most likely because a very long RNA molecule is probably not converted to cDNA as a full length molecule and sequencing capacity for molecules over 10 kb is quite low. As expected, probes 2 and 3 do not detect shorter isoforms but larger bands are visible. The antisense probe 2 should detect ORF64, if expressed. However, during these conditions the signal for ORF64 mRNA could not be detected. 18S and 28S ribosomal RNA were used as loading control as well as the integrity of the isolated total RNA. It should be noted that different probes work with different efficiencies for Northern blots and thickness of the bands should not be interpreted as a correlation of expression levels.

**Figure 2.**
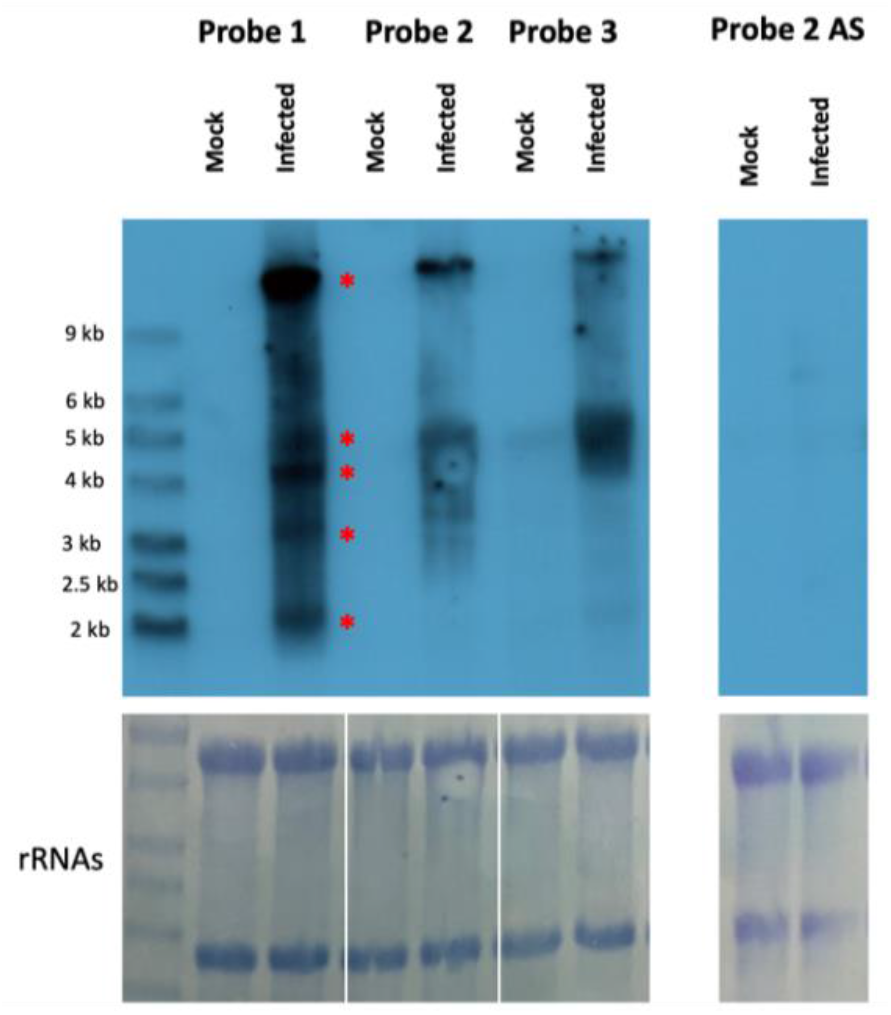
Northern blot for ORF6364 AS transcript with four different probes. The RNA ladder is shown and marked with the corresponding size. Kb kilobase. Red * indicates the transcripts with high scores discussed in the text.

### 3.3 Quantitative reverse transcription (qRT) PCR analysis

3 sets of primers with identical viral sequences to Northern blot probes were used for quantification of antisense RNA transcription. Initially, viral genome containing bacterial artificial chromosome (BAC) DNA is isolated from *E. coli* cells and the DNA is quantified using Nanodrop. By using DNA copy number calculation formula (DNA amount in ng x Avogadro’s number/ length of DNA × 10^9^ x 600), serial dilutions for 10^4^ to 10^7^ copy numbers were prepared and used as a template for qPCR primer efficiency. At around 10^6^ copy number the primers give Ct values lower than 20 suggesting good quality amplification (**Figure 3**). The primers are also used for the cDNA samples from mock or infected cells with or without reverse transcriptase enzyme. On average the Ct value ranges from 18.9 for set 2 to 21.1 for set 1. The primer efficiency depends on amplicon generation ability of the primer set and it is conceivable that each set would result in different Ct values. Usually, the best primer set is selected for further analysis however it should be noted that the best primer set selected here (set 2) depending on the Ct value will not be able to detect some of the transcripts from the region (**Figure 2**). Therefore it is essential to have a transcriptome map to start with, if quantification for a given lncRNA is studied.

**Figure 3.**
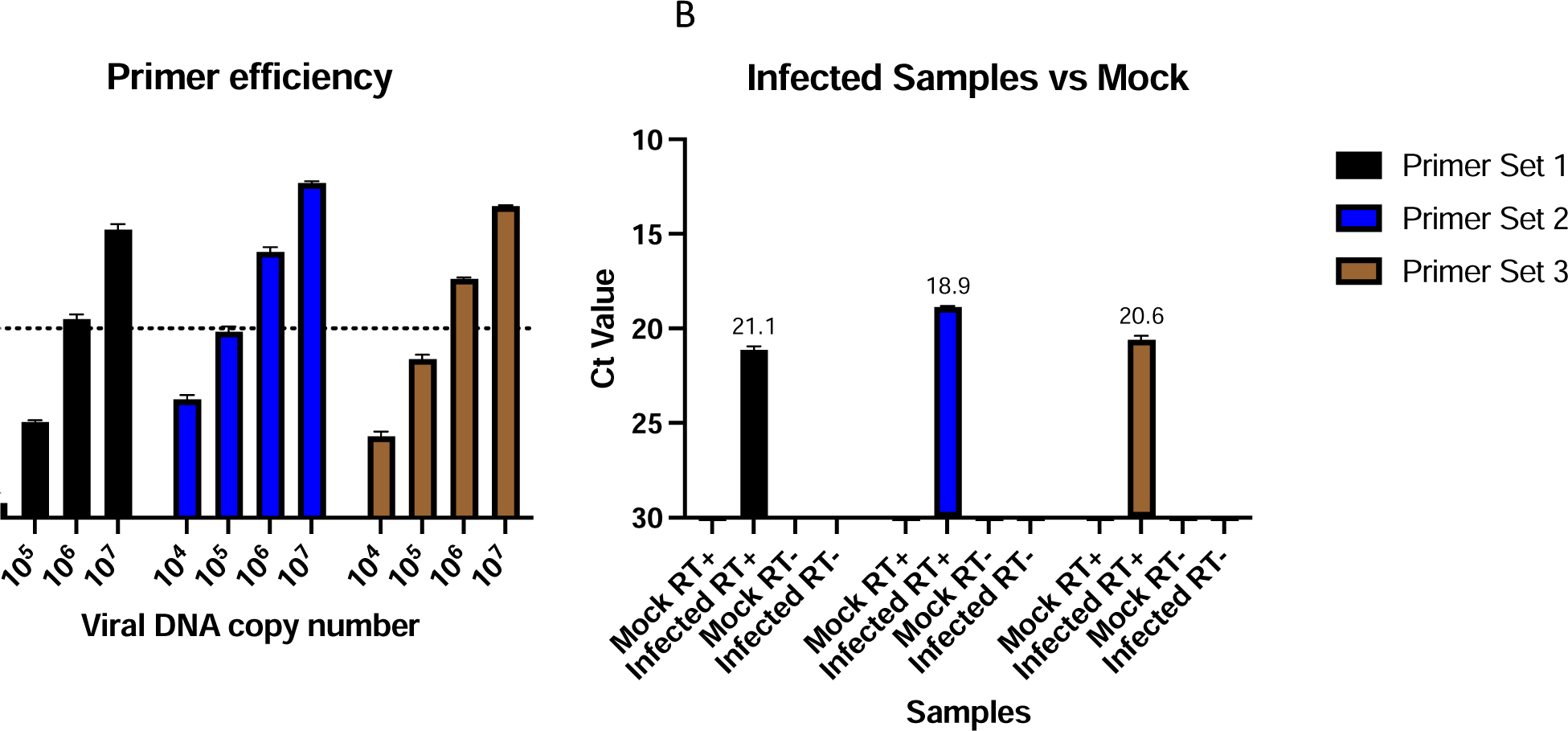
qPCR analysis for the ORF6364 antisense transcripts. The cycle threshold value (ct) is given in reverse order in the y axis.

### 3.4 Reverse transcription (RT)-PCR analysis

The splice junction is visible in some of the transcripts in **Figure 1**. In order to validate the splicing, RT PCRs are set for the mock or infected samples with set 2 forward and set 3 reverse primers. The full length amplicon is around 800bp (should be 778 bp) is visible from the positive control BAC DNA and infected RT+ samples. The spliced forms generate the lower molecular weight amplicon around 400 bp (should be 427 bp) which is consistent with the 351 nucleotide intron size. Even though the spliced isoforms are represented in low numbers with TRIMD analysis and are not detected with the Northern blots, it is clearly shown with a fainter band in the RT-PCR samples (**Figure 4A**). In order to grasp the full complexity of the transcripts from the locus, cDNA is prepared with the gene specific primer (**Table)** and PCR is performed with set 1 forward and set 3 reverse primers. The resulting amplicons were run on 10% polyacrylamide gel (**Figure 3B**). This clearly indicates the multiple variants can be detected by RT-PCR and viruses display much more complex transcriptional activity.

**Figure 4.**
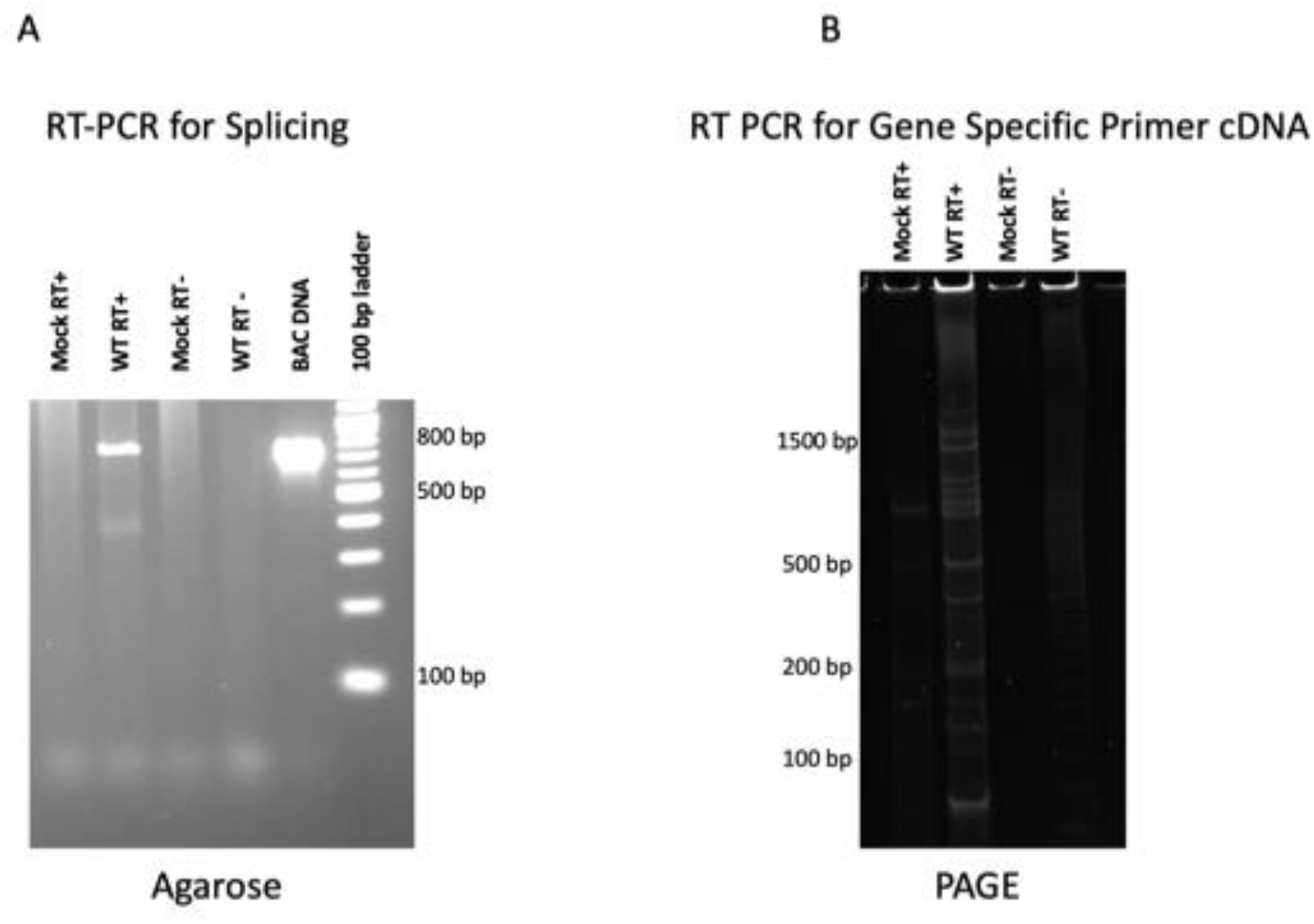
Reverse Transcription (RT)-PCR for the antisense transcripts in ORF63-64 locus. A) Set 2 forward and set 3 reverse primers are used and PCR products run on a 3% agarose gel. B) Set 1 forward and set 3 reverse primers are used and PCR products are loaded onto 10% PAGE gel.

### 3.5 Northern blot analysis for the time course and latency

Viruses utilize different transcriptional programs during infection depending on the time as well as state of the infection such as lytic or latency states. In order to examine whether transcripts differ through the course of infection, samples are collected at different time points after infection and Northern blot was performed. The earliest time point that any of the transcripts were detected was at 18 hours post infections and no apparent distinction regarding the isoforms was observed (**Figure 5A**). In addition, latency stage was tested. Since during latency much lesser viral transcripts are produced, the amount of RNA loaded on the gel from latently infected samples were increased to 15 μg while 3 μg of RNA was loaded from lytic sample as control (**Figure 5B**). During latency no specific transcripts were observed. However, upon induction of the reactivation with 12-O-tetradecanoylphorbol-13-acetate (TPA), a commonly used reagent to induce lytic replication for gammaherpesivuses, the transcripts start to become visible even though to a lesser extent.

**Figure 5.**
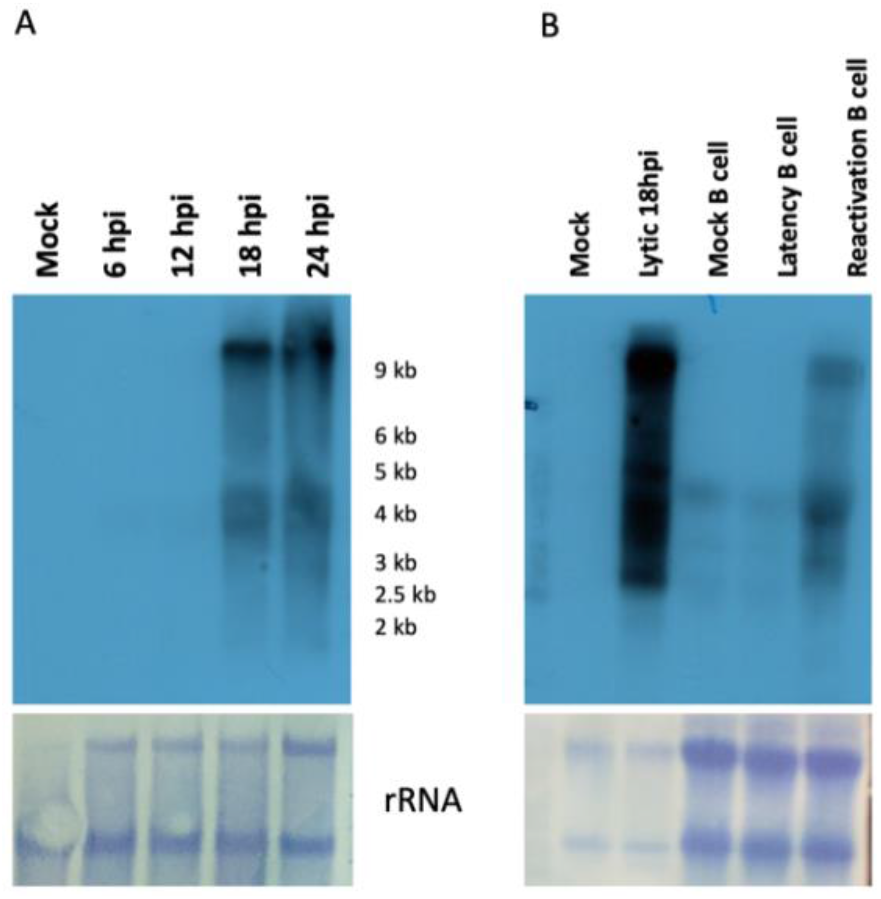
Northern blot for the time course (A) and latency (B)

## 4. Discussion and Conclusion

The work here investigates the ability of techniques used for lncRNA studies to grasp transcriptional complexity from a given locus. It has become clear in the field that genomes are pervasively transcribed and a vast pool of RNA products are generated (Derrien et al., 2012; Hangauer et al., 2013). One major method (if not only), to study the novel transcripts identified by the recently developed sequencing platforms, is the quantitative (real-time) PCR assay and majority of the published papers utilize this method for quantification of lncRNAs and mRNAs (Kolenda et al., 2021). This powerful method of quantification contains certain drawbacks which should not be overlooked, especially while working with gene dense large DNA viruses. The viral genomes contain many multigenic, antisense and read through transcripts (O’Grady et al., 2019; Tombácz et al., 2017). In order to understand the function of these newly identified class of RNA molecules, a detailed transcriptomic map of the genome proves to be useful to design primers as well as generation of mutants.

The locus discussed in this work was studied in several studies (Gredmark et al., 2007, p. 64; Sun et al., 2015; Xuan et al., 2013). From this locus, ORF64 protein, which is a large tegument protein with a deubiquinating protease activity, is produced. The work presented in these studies predates the most recent transcriptome analyses which revealed the antisense transcription. In some of these studies, data from qRT PCR assays are utilized to assess the expression level of ORF64. This proposes a question that how much of the expression is attributable to the ORF64 mRNA and how much of it is because of the antisense transcripts discussed here. Therefore, concluded from this work, it is simply suggested that a closer look to the region of interest through alternative RNA detection methods such as Northern blots alongside with the qPCR is essential, especially while working with viral transcripts.

## Acknowledgments/disclaimers/conflict of interest, if any

I would like to thank my PhD advisor Prof. Dr. Scott Tibbetts and Tibbetts Lab members. The laboratory work is performed at University of Florida during my postdoctoral research at Tibbetts Lab.

The work presented here did not receive any funding.

The author declares no conflict of interest.

